# Cerebellar theta and beta noninvasive stimulation rhythms differentially influence episodic memory versus language

**DOI:** 10.1101/2020.02.20.958397

**Authors:** Shruti Dave, Stephen VanHaerents, Joel L. Voss

## Abstract

The cerebellum is thought to interact with distributed brain networks to support cognitive abilities such as episodic memory and language. Episodic memory and language networks have been associated with distinct endogenous oscillatory activity frequency bands: theta (∼3-8 Hz) versus beta (∼13-30 Hz), respectively. We sought to test whether it is possible to toggle cerebellar participation in episodic memory versus language by noninvasively stimulating with theta versus beta rhythmic transcranial magnetic stimulation. Cerebellar theta stimulation improved episodic memory encoding but did not influence neural signals of language processing, whereas beta stimulation of the same cerebellar location increased neural signals of language processing but did not influence episodic memory encoding. This constitutes evidence for double-dissociation of cerebellar contributions to language versus episodic memory based on stimulation frequency pattern, supporting the hypothesis that the cerebellum can be biased to support these distinct cognitive abilities at the command of network-specific rhythmic activity.

## Introduction

The cerebellum contributes to a number of cognitive abilities, including episodic memory, language, decision-making, motor control, and others (Desmond and Fiez, 1998, Schmahmann, 2019). Many reports have particularly implicated lobules VI and VII, and especially Crus I and II (Stoodley and Schmahmann, 2010), in cognition. Portions of these cerebellar regions show resting-state fMRI connectivity and task-based co-activation with most major distributed cognitive brain networks (Buckner et al., 2011, Habas et al., 2009, King et al., 2019). For example, Crus I is a functionally heterogeneous lobule of posterolateral cerebellar cortex (Buckner et al., 2011, Strick et al., 2009, Stoodley, 2012) that interacts with the hippocampal network that supports episodic memory (Andreasen et al., 1999, Buckner et al., 2011, Fliessbach et al., 2007, Krienen and Buckner, 2009) and with the fronto-temporal-parietal network that supports language (De Smet et al., 2013, King et al., 2019, Pleger and Timmann, 2018), alongside other functional networks (Buckner et al., 2011, Habas et al., 2009). The goal of the current experiment was to evaluate whether such interactions are functional by testing whether noninvasive stimulation targeting cerebellar Crus I can differentially impact hallmark behavioral and neural correlates of episodic memory versus language.

The stimulation approach in this experiment was motivated by previous evidence that episodic memory and language networks are associated with distinct frequencies of oscillatory activity. Theta-band (∼3-8 Hz) interregional synchronicity of episodic memory network locations, including the cerebellum, has been associated with memory encoding and retrieval (Anderson et al., 2010, Lega et al., 2012, Rutishauser et al., 2010, Watrous et al., 2013, Hoffmann and Berry, 2009, Herweg et al., 2020), which is consistent with the dominant theta activity pattern of the hippocampus (Larson et al., 1986, Zhang and Jacobs, 2015, Bohbot et al., 2017). In contrast, beta-band (∼13-30 Hz) activity is prominent among language network locations including the cerebellum (Weiss and Mueller, 2012, Edagawa and Kawasaki, 2017, Bastiaansen and Hagoort, 2006), particularly during language tasks in which sentence context supports generating expectations or predictions for specific words (Engel et al., 2001, Lewis and Bastiaansen, 2015, Molinaro et al., 2016). We therefore hypothesized that transcranial magnetic stimulation applied to Crus I using theta versus beta rhythmic patterns could bias its participation in episodic memory versus language processing.

Several previous findings support the premise that noninvasive stimulation applied at theta versus beta frequencies may differentially affect episodic memory versus language networks, respectively. For instance, theta-burst transcranial magnetic stimulation (TMS) applied to a cortical location of the episodic memory network enhanced memory performance, whereas beta-frequency TMS of the same location did not affect memory (Hermiller et al., 2019). In contrast, beta-frequency TMS of a cortical location of the language network enhanced picture naming, whereas 1-Hz TMS did not alter picture naming (Mottaghy et al., 1999, Sparing et al., 2001). Importantly, these previous experiments used different stimulation locations within either the episodic memory or language network and did not assess the frequency-selectivity of stimulation effects on memory versus language. Therefore, it remains unclear if a single location such as Crus I can be biased to contribute to memory versus language by applying theta versus beta noninvasive stimulation. The experiment we conducted to test this possibility follows the logic of the “double-dissociation”, which is a powerful method for identifying functions supported by particular brain locations, networks, or processes (Teuber, 1955). This method involves demonstration that two distinct cognitive outcome measures are differentially modulated by two distinct lesions or interventions (Knowlton et al., 1996, Gil-Robles et al., 2013). The current experiment utilized this logic to test for rhythm-specific cerebellar stimulation effects on episodic memory encoding versus language measures.

We applied TMS targeting Crus I using either theta or beta rhythms and measured the impact of stimulation with a cognitive task that assessed highly canonical measures of language processing and episodic memory encoding (Figure 1). Stimulation effects on language were assessed via brain electrical activity measured during reading comprehension trials designed to elicit the event-related brain potential (ERP) N400 signal (Kutas and Federmeier, 2011, Kutas and Hillyard, 1980). Stimulation effects on episodic memory encoding were assessed by measuring subsequent high-confidence performance on a delayed recognition memory test (Eichenbaum et al., 2007, Shepard, 1967) administered after the period during which stimulation typically impacts neural function (Di Lazzaro et al., 2005, Huang and Kandel, 2005, Huang et al., 2005), and by examining ERPs recorded during encoding (Rugg and Yonelinas, 2003). As is typical for experiments following the double-dissociation logic, these measures of cognitive function differed fundamentally in format. Nonetheless, these distinct functional outcomes are advantageous because they are well-validated probes of language versus episodic memory (Rugg and Yonelinas, 2003, Yonelinas, 2001, Yonelinas, 2002, Kutas and Federmeier, 2011, Kutas and Hillyard, 1980), thus permitting strong inferences regarding any differential influence of cerebellar theta versus beta stimulation on cognition.

**Figure 1.**
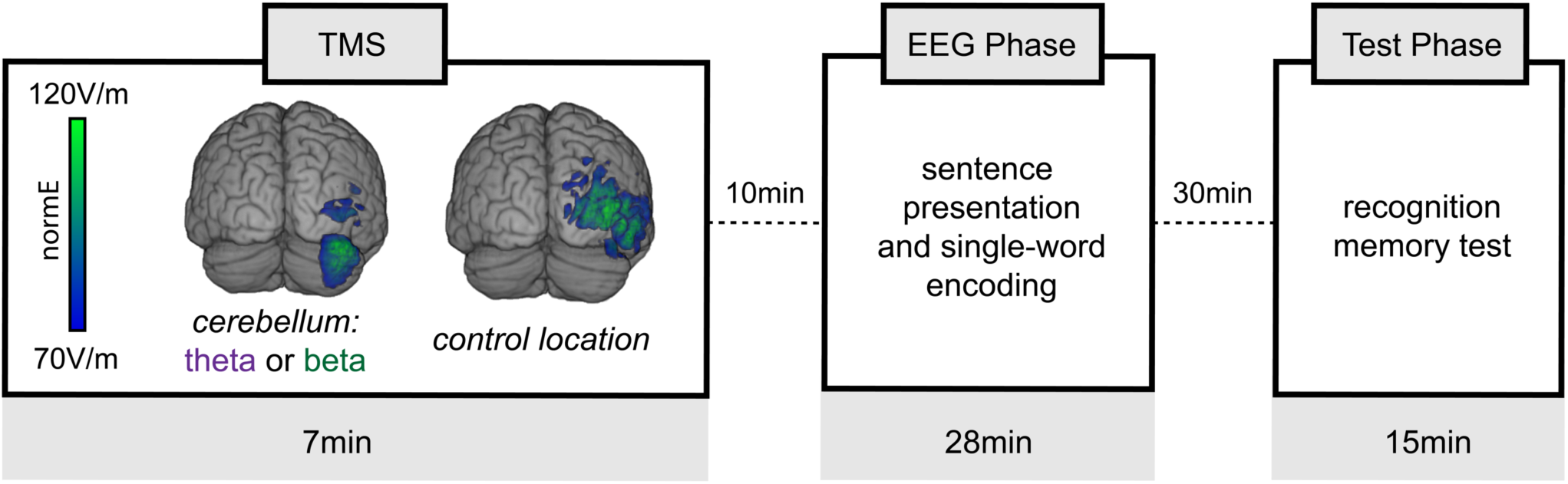
Experiment design. Each subject participated in three separate sessions. For each session, one of three TMS conditions was delivered: cerebellar theta (theta-burst), cerebellar beta (20 Hz), or control-location stimulation (either theta or beta). The induced electrical field (normE) is illustrated for a sample participant. Following stimulation on each session, subjects performed a mixed sentence reading and single-word encoding task while scalp EEG was recorded. This portion of the task occurred during the period when effects of stimulation on neural function were likely maximal. Recognition memory for words studied during the EEG phase was tested after a delay, such that retrieval occurred after the effects of stimulation had decayed (Methods).

We also applied the same stimulation rhythms to a control location of right lateral occipital cortex, as this location is unrelated to episodic memory and language networks and therefore served as an active-stimulation control (Figure 1). Each of the three stimulation conditions was delivered in a separate experimental session, using a within-subjects, counterbalanced design. We hypothesized that cerebellar beta stimulation would influence the N400 signal of linguistic prediction and not memory encoding, whereas cerebellar theta stimulation would influence memory encoding but not the N400. This hypothesized differential impact of stimulation rhythm on distinct language versus episodic memory assessments follows the same logic as aforementioned double-dissociation experiments, and would thereby provide strong evidence that lateral cerebellum (including Crus I) is rhythm-selective in its support of episodic memory versus language.

## Results

### Frequency-dependent stimulation effects on N400 correlates of language

We first tested whether cerebellar stimulation frequency influenced the N400 ERP component, which is robustly modulated by expectancy or predictability of words (Figure 2) (Kutas and Federmeier, 2011). The N400 is typically maximal for centro-posterior electrodes approx. 300-500ms following onset of less predictable as compared to highly predictable sentence-final words. As expected, subjects demonstrated typical effects of predictability on N400 amplitudes (i.e., ERP amplitude differences between high versus low predictability words measured at centro-posterior electrodes from 300-500 ms) for each of the three stimulation conditions (Figure 2; high versus low predictability amplitude for location-control stimulation: *t*(23) = 4.32, *p =* .0003, *d* = .83; cerebellar-beta stimulation: *t*(23) = 6.72, *p* < .0001, *d* = 1.33; cerebellar-theta stimulation: *t*(23) = 5.43, *p* < .0001, *d* = 1.16).

**Figure 2.**
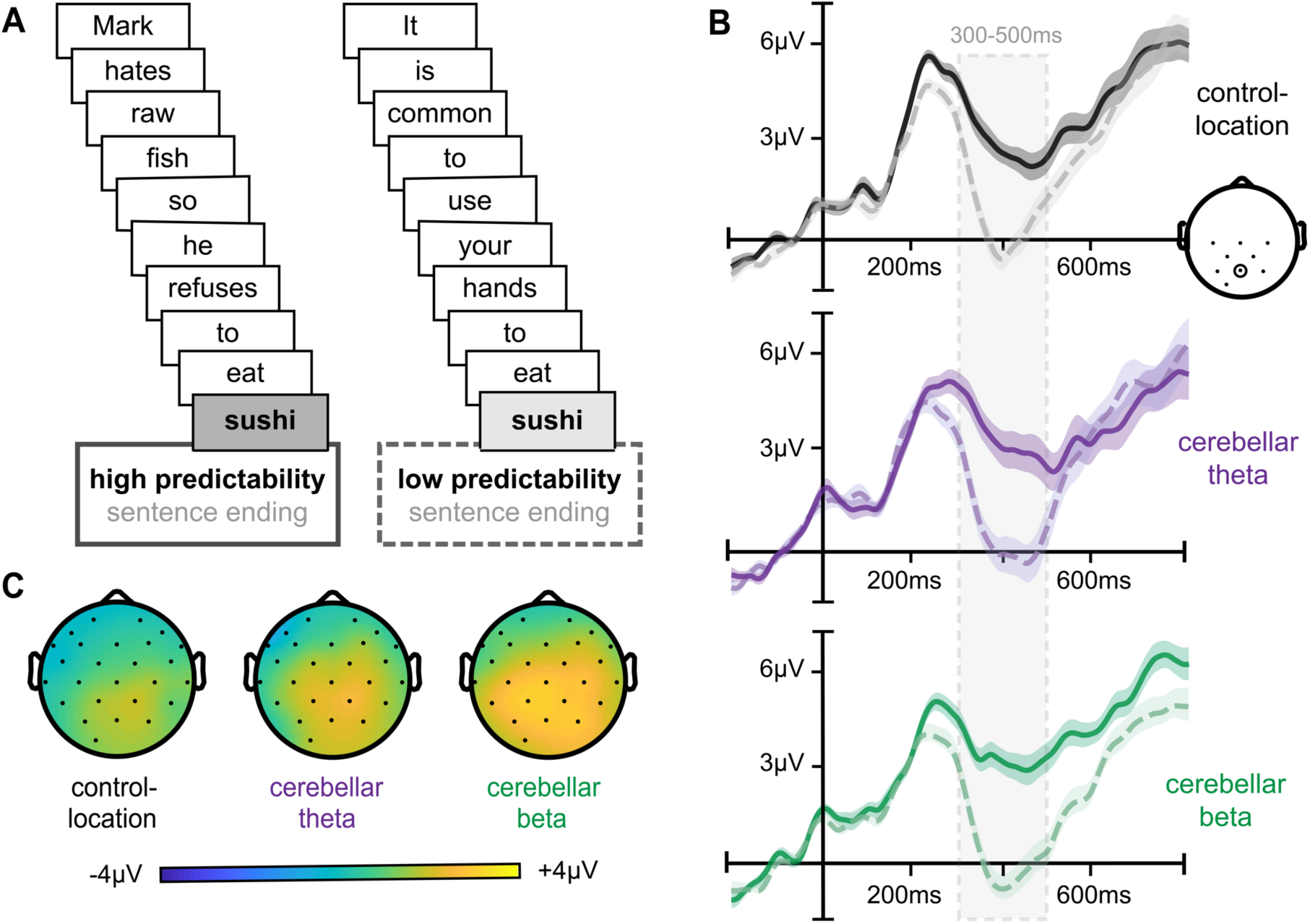
Typical ERP signals of language processing for each stimulation condition. (a) Example sentences for high versus low predictability sentence-final words. (b) Average ERPs to high (solid) and low (dashed) predictability words, plotted for a representative centro-posterior electrode (Pz) for all three stimulation conditions. (c) Scalp topography of the N400 effect (i.e., difference between high and low predictability words) for each stimulation condition shows a greater and more widespread N400 distribution following cerebellar beta stimulation relative to control-location stimulation.

As hypothesized, stimulation condition significantly influenced the effect of predictability on N400 amplitudes (Figure 3a; main effect: *F*(2,46) = 4.83, *p* = .01, *η*_*p*_ ^*2*^ = .21). Cerebellar beta stimulation enhanced the N400 effect amplitude relative to control-location stimulation (*t*(23) = 3.24, *p* = .004, *d* = .67). In contrast, cerebellar theta stimulation did not significantly influence the N400 predictability effect amplitude relative to control-location stimulation (*t*(23) = 1.45, *p* = .16) or relative to cerebellar beta stimulation (*t*(23) = 1.59, *p* = .13).

**Figure 3.**
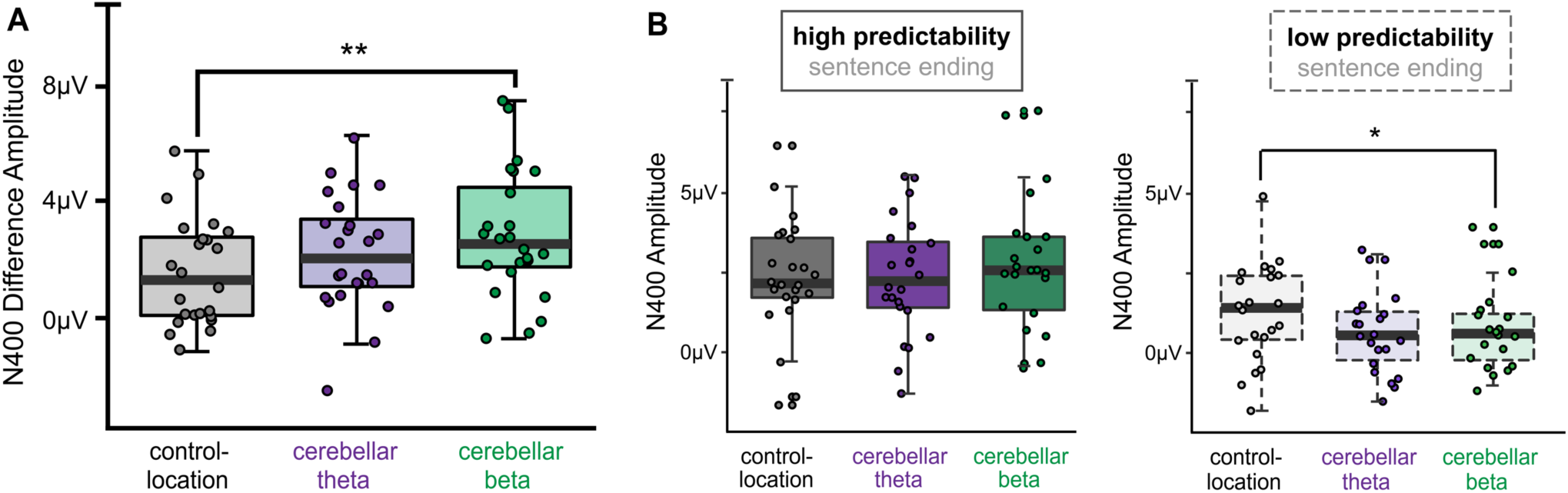
Cerebellar beta stimulation increased ERP signal of language processing. (a) Average N400 high minus low predictability difference amplitudes (300-500ms) for each stimulation condition. (b) Average N400 amplitudes for high predictability and low predictability words. Whiskers indicate first and third quartiles. Dots indicate values for individual subjects. ** *p* < .01. * *p* < .05.

As the N400 predictability effect was calculated as the difference between high and low predictability words (Kutas and Federmeier, 2011, Kuperberg and Jaeger, 2016), we performed post-hoc analyses to better understand how cerebellar beta stimulation increased the N400 effect relative to control-location stimulation (Figure 3b). Beta stimulation numerically but nonsignificantly increased N400 amplitudes to high predictability words relative to control-location stimulation (*t*(1, 23) = 1.27, *p* = .29). N400 amplitudes to low predictability words were significantly lower following cerebellar beta stimulation than control-location stimulation (*t(*1,23) = 4.84, *p* = .04, *d* = .17). Thus, the influence of beta stimulation on the effects of predictability on N400 reflected a combination of small increases in N400 amplitude to high predictability words and decreases in N400 to low predictability words, suggesting that word processing was more strongly influenced by the predictability of the sentence context (Kuperberg and Jaeger, 2016) following cerebellar beta stimulation.

Control analyses indicated that the effect of predictability on N400 difference amplitudes (i.e., difference between high and low predictability words) did not vary by session order irrespective of stimulation condition (all pairwise *p* values > 0.91). Thus, session-to-session practice or ordering effects did not significantly alter N400 amplitudes and therefore did not contribute to the influence of stimulation on N400 amplitudes. Furthermore, there was no significant difference in the effect of predictability on N400 amplitudes for subjects receiving beta stimulation of the control location versus subjects receiving theta stimulation of the control location (*t*(22) = 1.21, *p* = 0.24), indicating that rhythm-specific stimulation effects were unique to the cerebellum.

To evaluate cognitive specificity, we examined effects of stimulation on N400 amplitudes measured to words presented individually, as these words lacked sentence context and therefore should not have involved the same levels of language processing related to contextual predictability. N400 amplitudes to individually presented words did not vary significantly by stimulation condition (main effect: *F*(2, 46) = 1.09, *p* = 0.35, *η*_*p*_ ^*2*^ = 0.05; these ERPs are shown below: recognition memory ERPs). Therefore, the effects of cerebellar beta stimulation on N400 amplitudes were specific to sentence/contextual processing.

To further evaluate the selectivity of the predicted effect of beta stimulation on the N400, we performed exploratory control analyses of other ERP components often measured during language tasks. We examined the frontal post-N400 positivity (i.e., semantic P600, or PNP) associated with costs of incorrect predictions (Van Petten and Luka, 2012), the posterior post-N400 positivity (i.e., syntactic P600) associated with grammatical processing difficulties or animacy violations (Leckey and Federmeier, 2019), and the frontal N200/N250 associated with early visual word form processing (Holcomb and Grainger, 2006). There were no ERP amplitude differences between high and low predictability words for any of the three stimulation conditions for semantic or syntactic P600s (all pairwise *p* values > .08), but N200/N250 amplitudes differed by word predictability following cerebellar beta stimulation (*t*(23) = 3.09, *p* = .01, *d* = .13) and control-location stimulation (*t*(23) = 2.23, *p* = .04, *d* = .14), with a marginal difference for cerebellar theta stimulation (*t*(23) = 1.92, *p* = .07). However, contrary to the effects observed on the N400, stimulation condition did not modulate the effect of word predictability (high versus low predictability amplitude difference) for the N200/N250 component (*F*(2,46) = 0.33, *p* = .72). These control analyses therefore indicate that these non-hypothesized ERP components either were not sensitive to the task for any stimulation condition or showed no significant differential effect of stimulation rhythm.

### Rhythm-dependent stimulation effects on episodic memory encoding

We next evaluated effects of stimulation on episodic memory encoding of words presented individually, which were studied intermixed with sentence trials. These words were studied during the EEG phase within 45 minutes of stimulation delivery, i.e., within the period when the stimulation parameters have been found to impact neural activity and cognitive function (Hoogendam et al., 2010, Rossi et al., 2009, Thut and Pascual-Leone, 2010). The recognition memory test was administered following a delay (Figure 1), after the expected duration of impact for the current stimulation conditions. During the recognition test, subjects discriminated studied words from novel foils and simultaneously rated memory confidence (Jacoby, 1983, Kelley and Wixted, 2001, Yonelinas, 1999). For all stimulation conditions on average, subjects made more confident “old” responses to studied words than novel foils (Table 1; *t*(71) = 15.17, *p* < 0.001, *d* = 2.52; d′ *M* = 1.65, d′ *SD* = 0.82), indicating successful discrimination. However, discrimination was at chance when subjects made guess responses (similar “old” endorsement rates for studied words and novel foils; *t*(71) = 0.60, *p* = 0.55; d′ *M* = 0.02, d′ *SD* = 0.42). As intended, only confident recognition trials were highly accurate and likely reflected the recollection component of episodic memory that depends on the hippocampal network (Eichenbaum et al., 2007, Diana et al., 2007, Yonelinas, 2002, Yonelinas et al., 2005). We therefore tested the effects of cerebellar stimulation on the accuracy of confident responses.

**Table 1.** Response rates to old and new words for confident and guess responses. Values are means across subjects. Parentheses indicate standard deviation of the mean. Significant pairwise comparisons are described in text.

|  |  |  | Response type |  |  |  |
| --- | --- | --- | --- | --- | --- | --- |
|  |  |  | Old |  | New |  |
|  |  |  | Confident | Guess | Confident | Guess |
| Stimulus type | Old words | Control-location | 0.45 (0.24) | 0.16 (0.06) | 0.15 (0.10) | 0.07 (0.06) |
|  |  | Cerebellar theta | 0.50 (0.21) | 0.16 (0.13) | 0.15 (0.10) | 0.04 (0.04) |
|  |  | Cerebellar beta | 0.46 (0.23) | 0.14 (0.09) | 0.15 (0.11) | 0.07 (0.05) |
|  | New words | Control-location | 0.19 (0.22) | 0.20 (0.19) | 0.28 (0.24) | 0.51 (0.28) |
|  |  | Cerebellar theta | 0.16 (0.19) | 0.19 (0.18) | 0.29 (0.21) | 0.52 (0.28) |
|  |  | Cerebellar beta | 0.16 (0.19) | 0.23 (0.16) | 0.24 (0.17) | 0.55 (0.25) |

Confident recognition discrimination accuracy was influenced by stimulation condition (Figure 4; main effect: *F*(2, 46) = 5.61, *p* = 0.01, *η*_*p*_ ^*2*^ = 0.24). Cerebellar theta stimulation enhanced accuracy relative to cerebellar beta stimulation (*t*(23) = 2.34, *p* = 0.03, *d* = 0.35) and relative to control-location stimulation (*t*(23) = 3.84, *p* = 0.001, *d* = 0.38). In contrast, cerebellar beta stimulation did not significantly influence accuracy relative to control-location stimulation (*t*(23) = 0.06, *p* = 0.95). The beneficial effects of cerebellar theta stimulation on discrimination accuracy were related to increased hit rates relative to control-location stimulation (*t*(23) = 2.09, *p* = 0.05, *d* = 0.20) as well as reduced false alarm rates relative to control-location (*t*(23) = 2.26, *p* = 0.03, *d* = 0.27) and cerebellar beta stimulation (*t*(23) = 2.91, *p* = 0.01, *d* = 0.57) (Table 1).

**Figure 4.**
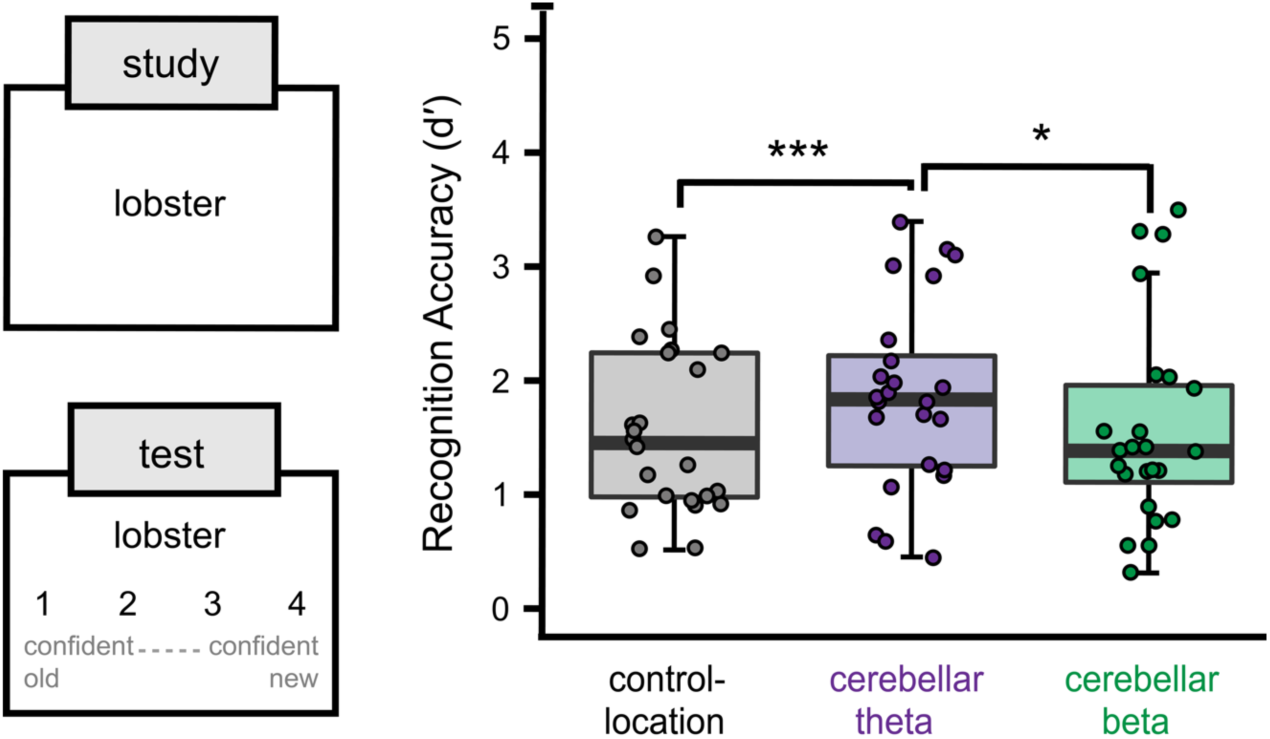
Cerebellar theta stimulation enhances episodic memory. Words were studied individually during the EEG phase and a recognition memory test was administered after a delay. Subjects attempted to discriminate studied words from novel foils and simultaneously rated confidence. Recognition discrimination accuracy for confident responses (d′) varied by stimulation condition, as shown via box-and-whisker plots with whiskers marking first and third quartiles. Dots indicate values for individual subjects. * *p* < .05. *** *p* < .001.

Control analyses indicated that recognition discrimination accuracy did not vary by session order irrespective of stimulation condition (all pairwise *p* values > 0.14). Thus, session-to-session practice or ordering effects did not significantly contribute to the influence of stimulation on memory. Furthermore, there was no significant difference in accuracy for subjects receiving beta versus theta stimulation for the control-location condition (*t*(22) = 0.79, *p* = 0.46), indicating that rhythm-specific stimulation effects were unique to the cerebellum.

We performed two further analyses to support the assumption that these stimulation effects were relatively specific to episodic memory encoding (as the encoding period immediately followed stimulation and the recognition test was administered at a delay that is typically considered to be past the duration of stimulation effects on cognition for the current parameters (Hoogendam et al., 2010, Rossi et al., 2009, Thut and Pascual-Leone, 2010)). If the effects of stimulation persisted during the recognition test, they would be expected to decay over the course of the recognition test, leading to variable performance between early versus late memory testing, which might vary as a function of stimulation condition. We therefore tested whether theta versus beta stimulation conditions differentially influenced performance for the first and last third of memory test trials. Confident recognition discrimination accuracy did not vary between early and late memory trials for either stimulation condition (cerebellar beta stimulation: *t*(1, 23) = 0.83, *p* = .41; cerebellar theta stimulation: *t*(1, 23) = 0.29, *p* = .77), with no differential early versus late difference by condition (*t*(1,23) = 0.45, *p* = .66). This finding supports the conclusion that the impact of cerebellar theta stimulation on memory reflected an influence on encoding.

Next, we analyzed effects of stimulation on ERPs measured during single-word encoding, as an influence of cerebellar stimulation on these ERPs would further suggest an impact of theta stimulation on encoding. We focused on late-positive ERP amplitudes, as this signal has been associated with memory formation in many experiments (Rugg and Yonelinas, 2003, Voss and Paller, 2007). To ensure roughly equal trials across stimulation conditions, we analyzed late-positive ERP amplitude for all single-word encoding trials irrespective of subsequent recognition performance (Figure 5). Amplitudes were weakly modulated by stimulation condition (*F*(2, 46) = 2.96, *p* = .06, *η*_*p*_ ^*2*^ = .13), reflecting relatively lower late-positive amplitude following cerebellar theta stimulation versus location-control stimulation (*t*(23) = 2.06, *p* = .05, *d* = .56). ERP amplitudes following cerebellar beta stimulation did not significantly differ from either cerebellar theta (*t*(23) = 1.20, *p* = .24) or location-control stimulation (*t*(23) = 1.60, *p* = .12). Thus, cerebellar theta stimulation weakly influenced ERP signals typical of word encoding when measured across all trials. Control analyses indicated that session order did not influence late-positive ERP amplitudes (all pairwise *p* values > 0.12). Additionally, subjects receiving beta versus theta stimulation of the control location did not have significantly different ERP amplitudes (*t*(22) = 1.84, *p* = 0.08). These results indicate that stimulation effects on ERP signals associated with encoding were specific to theta delivered to the cerebellum even when calculated on all words in aggregate, without considering trial-specific memory outcomes.

**Figure 5.**
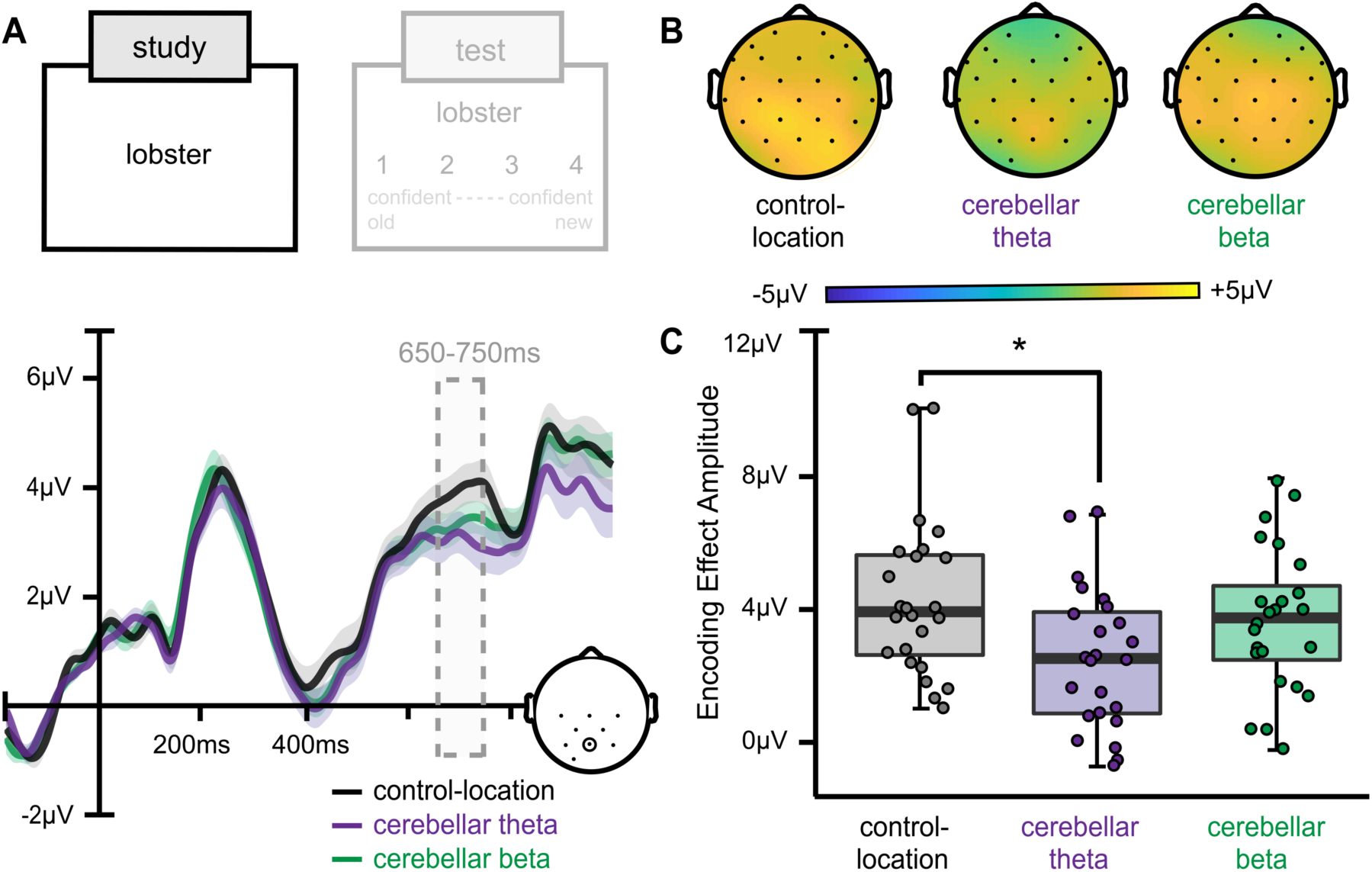
Cerebellar theta stimulation effects on ERP correlates of memory encoding. (a) Average ERPs during single-word presentation are plotted for a representative centro-posterior electrode (Pz). Encoding effect amplitudes (650-750ms) differed by stimulation condition, with cerebellar theta stimulation reducing amplitudes relative to control-location stimulation. (b) Scalp topography of the parietal memory effect for each stimulation condition shows a greater, more widespread positive distribution following control-location stimulation relative to cerebellar theta-burst stimulation. (c) Box-and-whisker plots of mean amplitudes between 650-750ms per stimulation condition. Whiskers indicate first and third quartiles. Dots indicate values for individual subjects. Shading on ERP waveforms indicates standard deviation of the group mean. * *p* < .05.

### Effects of stimulation on language versus episodic memory

Beta versus theta cerebellar stimulation had opposite effects on language versus episodic memory. This pattern can be appreciated by considering standardized effect sizes. Compared to the control-location stimulation condition, cerebellar beta stimulation had a larger impact on the N400 signal of language than theta stimulation, whereas the opposite pattern (greater impact of theta than beta) was identified for high-confidence recognition memory accuracy and late-positive ERPs during encoding (Figure 6). This supports the conclusion that language and memory were doubly dissociated based on different effects of beta versus theta cerebellar stimulation, respectively.

**Figure 6.**
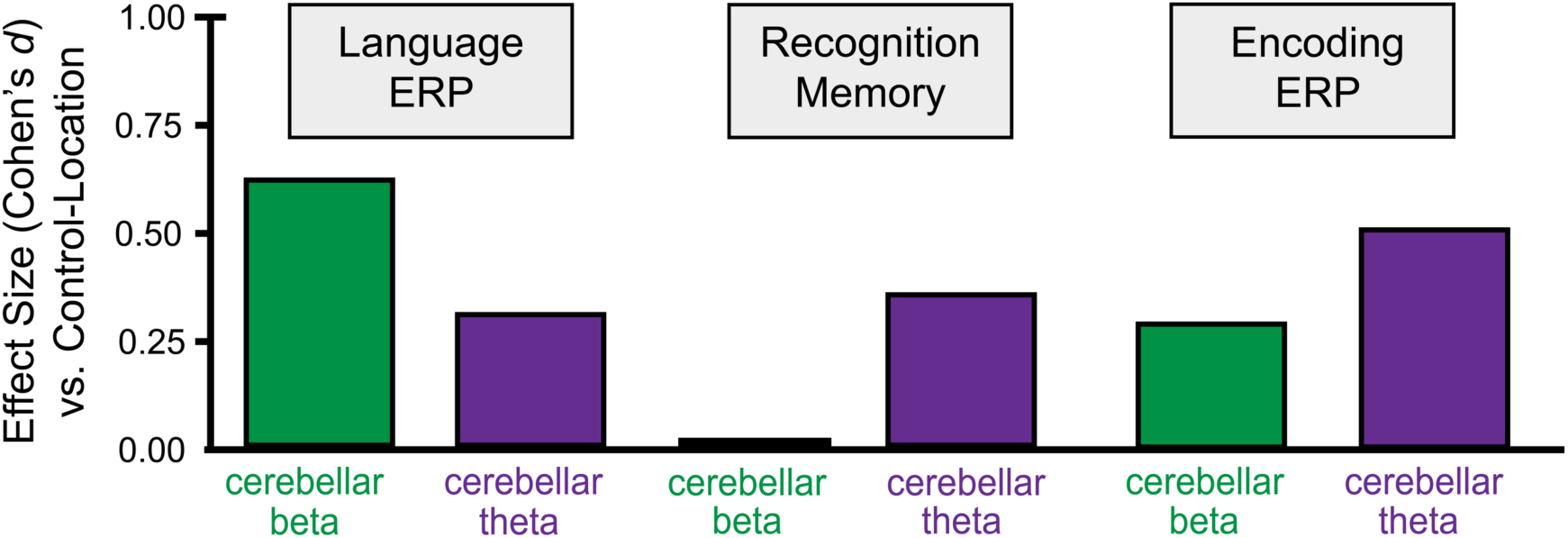
Opposite effects of stimulation rhythms on language versus episodic memory. Standardized effect sizes for cerebellar theta and beta compared to control-location stimulation for N400 amplitudes, confident recognition memory accuracy (d′), and encoding-related ERPs.

## Discussion

Language processing and episodic memory encoding were differentially affected by cerebellar TMS applied to mimic endogenous activity rhythms of their respective networks. The nature of the effect of cerebellar beta stimulation on N400 ERP amplitudes suggests enhancement of language function, as predictable words elicited greater N400 amplitude whereas unpredictable words elicited reduced N400s. These findings indicate that cerebellar beta stimulation enhanced discrimination of contextual predictability. In contrast, cerebellar theta stimulation increased recognition discrimination accuracy, and this effect was likely driven by stimulation-related changes in encoding. In sum, our results suggest cerebellar beta and theta stimulation had dissociable effects on language versus episodic memory, respectively. This differential effect on cognitive functions by stimulation rhythms follows the logic of the neuropsychological double dissociation (Teuber, 1955, Dunn and Kirsner, 2003), suggesting that cerebellar involvement in these functions is dissociable under the command of rhythmic stimulation. It has been suggested that cerebellum might coordinate with distributed networks via interregional synchronicity of oscillatory activity (Mantini et al., 2007, Uhlhaas et al., 2009), as neural oscillations can support temporal coordination of activity across separated brain regions in support of cognitive processing (Buzsaki and Draguhn, 2004, Engel and Fries, 2010). Thus, our findings of rhythm-specific effects of brain stimulation on distinct cognitive outcomes are consistent with this “communication-by-synchrony” hypothesis.

The distributed network associated with the hippocampus has long been identified as important for episodic memory (Mesulam, 1990, Eichenbaum et al., 2007, Ranganath and Ritchey, 2012), and network interactions associated with memory performance have been correlated to synchronous activity in the theta frequency band (Lega et al., 2012, Anderson et al., 2010). Recent evidence has demonstrated resting-state fMRI connectivity between the hippocampus and the ventrolateral cerebellum (Buckner et al., 2011) as well as theta-band phase synchrony between the hippocampus and cerebellum during memory-dependent tasks (Wikgren et al., 2010). A previous experiment found that theta TMS targeting Crus I/II alters resting-state fMRI connectivity between the cerebellum and the hippocampal network as well as within the hippocampal network (Halko et al., 2014). Our results provide a cognitive corollary to this effect. We further show theta was more effective than beta stimulation for memory enhancement, which is consistent with a previous study using parietal cortex TMS (Hermiller et al., 2019), and suggests that theta is relatively privileged in its ability to alter function of the hippocampal network.

The N400 is an exceptionally well-characterized neural index of semantic processing, and thereby a very useful functional outcome of the language network (Kutas and Federmeier, 2011, Kutas and Hillyard, 1980). N400 differences between words that are highly predictable versus less predictable has been interpreted in two primary ways: as reflecting differences between the effort needed to semantically (i.e., contextually) integrate the incoming word into the ongoing sentence, or as reflecting differences in how sentence context can boost prediction or pre-activation of a specific word (Kuperberg and Jaeger, 2016). Although our experimental design is agnostic to this debate, it is noteworthy that effects of cerebellar beta stimulation on the N400 were primarily driven by changes to the processing of low-predictability sentence-final words. Stimulation may have influenced how words that were unlikely to be predicted or pre-activated were processed (or, alternately, may have affected processing of words that require more effect to contextually integrate into the ongoing sentence). Importantly, effects of cerebellar stimulation on the N400 were specific to beta. In EEG and MEG studies examining oscillatory activity during sentences that typically elicit N400 effects, beta power was altered in response to contextual predictability (Wang et al., 2012, Luo et al., 2010, Molinaro et al., 2016). Whether these changes in beta are synchronous across regions in the frontal-temporal-parietal network that supports language comprehension (Hagoort, 2014, Lau et al., 2008) is still unknown. Nonetheless, our findings suggest that beta is relatively privileged in its ability to alter the language functions characterized by the N400.

The current “double dissociation” based on stimulation rhythm is notable with respect to the possibility of shared functional neuroanatomy for language processing and episodic memory. Some have hypothesized hippocampal involvement in lexical prediction (Bonhage et al., 2015, Mullally and Maguire, 2014, Davachi and DuBrow, 2015, Buckner, 2010), perhaps due to the role of hippocampus in semantic memory (Piai et al., 2016) and/or in domain-general processing of prediction error ((Covington and Duff, 2016); but see (Ryskin et al., 2019)). Consistent with this hypothesis, cerebellar theta stimulation influenced both functions but with greater effects on episodic memory, whereas cerebellar beta stimulation exclusively affected language processing and not episodic memory (Figures 3 and 5). Nevertheless, we identified differential patterns of influence consistent with the functional dissociation of language versus episodic memory based on the rhythm of cerebellar stimulation, indicating dissociation of both these functions and cerebellar contributions to them.

Our results are consistent with theories that brain stimulation impacts networks by entraining their oscillatory activity (Kim et al., 2016, Lea-Carnall et al., 2017, Buzsaki, 2002, Lakatos et al., 2019). Notably, we do not suggest that network activity remained entrained to the applied stimulation rhythm long after stimulation delivery to achieve the current results, but rather that a lasting impact on network function was achieved by rhythm-matched stimulation due to an entrainment mechanism. In contrast to these resonance-oriented theories, compensation-oriented accounts propose that stimulation effects on local brain activity result in the opposite effect on connected regions, via an unknown homeostatic mechanism (Cocchi et al., 2015, Eldaief et al., 2011, Steel et al., 2016). The continuous theta-burst stimulation parameters used in the current theta condition has been characterized as locally inhibitory whereas beta (20-Hz) stimulation has been characterized as locally excitatory, based almost entirely on their effects when applied to primary motor cortex (Chen et al., 2003, Di Lazzaro et al., 2005, Pascual-Leone et al., 1994). Compensation-oriented theories would therefore predict opposite effects of stimulation on cognitive processing (facilitation versus disruption). Our findings are inconsistent with this prediction, as stimulation frequencies led to different levels of enhancement for different functions, but not to opposite effects. Likewise, Hermiller et al. (2019) found that continuous theta-burst stimulation of a parietal cortex location of the episodic memory network increased episodic memory and memory-related fMRI connectivity to a greater degree than did beta stimulation, but beta stimulation did not result in an opposite effect relative to sham stimulation. It has also recently been demonstrated that stimulation parameters have non-uniform effects when applied to different cortical areas (Castrillon et al., 2020), calling into question the simple excitation/inhibition dichotomies derived from studies exclusively of primary motor cortex. Collectively, these results are consistent with resonance-oriented accounts of brain stimulation effects on cognition, whereby stimulation patterns matched to network-specific oscillations optimally facilitate network function. Nonetheless, long-lasting impact of effective rhythms are governed by yet-unknown mechanisms, which could include a variety of neuroplasticity-related processes and are unlikely to reflect long-lasting resonance of neural activity following stimulation.

One limitation of the current study is that mechanistic interpretations are limited by the nature of the episodic memory and language outcomes measured. It is thought that the cerebellum controls the intricate timing of neural processing throughout distributed cortical networks in a frequency-specific manner (McAfee et al., 2019). There are compelling models for the relevance of theta-gamma coordination for episodic memory (Shirvalkar et al., 2010, Lisman and Jensen, 2013, Buzsaki and Schomburg, 2015) and of beta coordination with other frequencies for linguistic prediction (Arnal et al., 2015, Arnal and Giraud, 2012, Saleh et al., 2010). Our noninvasive measurement was not able to determine whether stimulation altered circuit-specific timing of activity, and therefore could not evaluate such mechanisms. However, it is notable that the frequency-specific impact of stimulation on cognitive function is consistent with the idea that oscillatory activity was affected by cerebellar stimulation, at least temporarily. Further, although the experiment design and results suggest that the effect of cerebellar theta stimulation was on memory encoding, we did not also test for selective encoding effects by delivering stimulation between encoding and retrieval, and so cannot be entirely confident that only encoding was improved versus other stages of memory processing. Finally, although we targeted cerebellar Crus I, the induced electrical field could have influenced other areas of cerebellum, and our conclusions regarding the functional neuroanatomical specificity of the observed influences of stimulation are limited by the lack of localization data, such as provided by fMRI or related methods. These are issues that could be addressed in future experiments.

The current findings could be relevant to the use of noninvasive brain stimulation for the treatment of language and episodic memory impairments, such as those that occur due to neurodegeneration of corresponding language and episodic memory networks (Golomb et al., 1993, Mesulam et al., 2014). Rhythm-specific stimulation effects on language versus episodic memory could provide a means to efficiently target specific symptoms in specific patient groups to achieve relatively personalized treatments. Furthermore, the cerebellum may be superior to cerebral cortical targets for stimulation in neurodegenerative disease because it is relatively less affected by neuropathological processes (Marcyniuk et al., 1986, Mann et al., 1990), although it is not immune to them (Guo et al., 2016, Schmahmann, 2016). Further investigation of rhythm-specific effects of cerebellar stimulation could thus advance understanding of cerebellar contributions to cognitive function while also motivating novel treatment approaches for cognitive impairment.

## Methods

### Participants

Subjects (N = 25) had normal or corrected-to-normal vision and passed MRI and TMS safety screenings (Rossi et al., 2009) supervised by S.V. Subjects had no known history of psychiatric or neurological disorder. All provided written informed consent to a protocol approved by the Institutional Review Board at Northwestern University and were paid for their participation. Data from one subject were excluded from analyses due to excessive artifacts in EEG recordings (see below). Reported analyses thus include 24 adults (16 women; average age: 25.3 years; range: 18-37 years).

### Experiment design

Prior to the three experiment sessions, each subject completed a structural MRI scan to provide anatomical localization for TMS targeting. The experiment used a within-subjects design across three sessions performed on non-consecutive days (average inter-session interval: 6 days; range: 2-26 days). During each session, one of three stimulation conditions was delivered immediately prior to the EEG phase, which was then followed by the test phase (Figure 1). The EEG phase comprised approximately 28 minutes during which sentence and single-word materials were presented while EEG was recorded (see below). Stimulation order across sessions was counterbalanced across subjects. Visual stimuli were counterbalanced across stimulation conditions and session orders. EEG caps were affixed to subjects and prepared prior to stimulation, such that presentation of experimental materials began 10 minutes after the final stimulation pulse for each session. The EEG phase was followed by a 30-minute delayed word recognition task test phase, during which EEG was not recorded.

### TMS protocol

During each session, subjects first received one of three TMS conditions: cerebellar theta (5 Hz delivery of 50-Hz bursts), cerebellar beta (20 Hz), or location-control stimulation. Location-control stimulation was delivered at the right lateral occipital cortex and was counterbalanced between theta and beta pattered stimulation across subjects.

MRI data were collected using a Siemens 3T Prisma whole-body scanner with a 64-channel head-neck coil located in the Northwestern University Center for Translational Imaging (CTI). Structural images were acquired using a T1-weight MPRAGE sequence (176 frames; TE: 1.69ms; TR: 2170ms; TI: 1160ms; flip angle: 7°; voxel resolution: 1.0 × 1.0 × 1.0 mm; 1-mm thick sagittal slices; 256 × 256mm FOV; scan duration: 5.1min). Structural MRI data were preprocessed using AFNI software (Version AFNI_19.3.04) (Cox, 1996). Scans were skull stripped (*3dSkullStrip*) and warped into Talairach-Tournoux (TT) stereotactic space using the TT_N27 atlas (*auto_tlrc*). Lateral-cerebellar and occipital control-location stimulation sites were selected using *a priori* coordinates and are listed in MNI space for convenience (cerebellum MNI: 41, −79, −39 (cf. (Halko et al., 2014)); occipital MNI: 47, −78, −13 defined based on lack of membership in memory and language cortical network parcellations (Yeo et al., 2011, Shain, 2019)). Each stimulation location was adjusted per subject to fall on the nearest adjacent brain surface (mean distance from default target: 1.7 mm for cerebellum, 3.6 mm for occipital) and then transformed back into subject-specific original MRI space for anatomically guided TMS. The average distance between cerebellar and control-location stimulation targets in original space was 29.6 mm (*SD* = 1.7 mm).

Beta and theta stimulation both comprised 600 bipolar pulses applied at individualized intensity. For beta stimulation, pulses were delivered in fifteen 40-pulse 20-Hz trains (2sec on, 28sec off; approx. 7 min. total duration; i.e., 20-Hz repetitive TMS). For theta stimulation, 50-Hz pulse triplets were delivered every 200ms (5 Hz) for 40 seconds continuously (i.e., continuous theta-burst stimulation). To better match the subjective duration of stimulation conditions, theta stimulation was preceded by approx. 6.5 minutes of beta stimulation delivered at a very low, sub-threshold intensity (10% of resting motor threshold). Each subject received theta stimulation and beta stimulation of the cerebellar target across different sessions. For the control-location session, half of the subjects received theta stimulation and half received beta stimulation. All stimulation conditions were counterbalanced across session order and were matched across sessions for stimulation intensity (calibrated per subject). Subjects were informed that stimulation pattern and location and thus sensation would vary among sessions but were unaware of location- or stimulation-specific study hypotheses.

We delivered TMS via a MagPro X100 system with a Cool-B65 coil (MagVenture A/S, Denmark) using frameless stereotactic guidance (Localite GmBH, Germany). Maximum stimulation intensity was calibrated to 80% of the resting motor threshold identified for the right *abductor pollicis brevis* before the experimental sessions. Intensity was adjusted for comfort in 13 subjects, resulting in an average stimulation intensity of 78.7% resting motor threshold (42.8% stimulator maximum output, range: 32-55%). For subjects with adjusted intensity, the same intensity was used for all stimulation conditions.

TMS was delivered to optimize induced electric field orientation, directed medial-rostrally for the cerebellum and medial-laterally for the occipital cortex (Janssen et al., 2015). Finite element modeling using subject-specific MRI was used to calculate induced electrical field following cerebellar and location-control stimulation (as shown for a representative subject in Figure 1), implemented in SimNIBS version 3.0 (Thielscher et al., 2015).

### Combined prediction and episodic memory task

*EEG phase.* 80 sentences and 64 single words were presented during each session. Subjects were instructed to read each sentence for comprehension, which was tested by manual response to true-false statements following 16 trials. Subjects were additionally informed that they would be tested on word memory. At each session, subject performed a brief (approximately 5min) practice including the delayed memory test.

Stimuli included 480 sentence pairs with 240 unique sentence-final critical words (similar to materials presented in (Dave et al., 2018)). Two factors varied between sentences in each pair: *cloze*, or the likelihood a sentence would end in a specific sentence-final word, and *contextual constraint*, or the highest probability a sentence would end in any final word. One sentence of each pair was strongly constrained towards a highly expected (i.e., high cloze) ending (average cloze: 90%; range: 80-100%), e.g., *Mark hates raw fish, so he refuses to eat <u>sushi</u>*. These endings are referred to as “high predictability”. The second sentence in each pair was weakly constrained (average constraint: 21.3%; range: 6-30%) and ended with a low cloze, or unlikely to be predicted, final word (average cloze: 1.1%, range: 0-4%), e.g., *It is common to use your hands to eat <u>sushi</u>*. These endings are referred to as “low predictability.” Averages for cloze and constraint were obtained from 50 additional adults (average age: 36.4 years; range: 22-68 years) recruited and compensated via Amazon Mechanical Turk.

Sentence length was approximately matched across sentence pairs generated for the prediction task (i.e., within 2 words for each pair; average length: 9 words; range: 6-11 words). Critical words had a mean SUBTLEX-US frequency count of 1215, mean length of 5.4 letters, and a mean concreteness rating of 3.9 (Brysbaert et al., 2014)). Frequency, word length, and concreteness ratings of sentence-final words were matched across the 80 sentences selected for each session.

We created a list of 192 separate words for single-word trial presentation. For the single words, the mean frequency count was 1182, mean word length was 5.4 letters, and a mean concreteness rating was 3.8. 64 words were presented during each session, matched on frequency, length, and concreteness across sessions.

During the EEG phase of each session, each subject read 40 high predictability and 40 low predictability sentences that were randomly intermixed, such that each critical word and its corresponding sentence was presented once per subject. The sentences were counterbalanced across two experimental lists so that each target word appeared equally often in high and low predictability sentences. To test sentence comprehension, subjects were asked to evaluate true-false statements on 20 percent of trials (average accuracy: 92.3%, range: 75-100%), indicating that subjects both understood sentence content and were paying adequate attention while reading. 64 single-word trials were presented during each session, randomly intermixed with sentence trials.

Prior to each trial, a cue (e.g., SENTENCE or WORD) was displayed for 2000ms to indicate upcoming trial type to subjects. Cue presentation was followed by a 4000ms display of a central fixation cross. Sentences were presented via rapid serial visual presentation (RSVP) with a stimulus-onset asynchrony of 600ms (300ms/word). Single words were similarly presented for 300ms following two buffer items (“XXXXXX”) used to standardize reading preparedness between sentence and single-word trials. 1200ms of blank screen were shown after sentence-final target words and single-word trials, followed by a 4000ms display of a central fixation cross between trials.

Continuous EEG was recorded from 27 active Ag/AgCl electrodes mounted in an elastic cap (Brain Vision LLC, actiCAP). Electrodes were not positioned at areas where TMS was delivered in order to facilitate cerebellar and occipital stimulation. To monitor saccades and blink activity, four additional electrodes were placed below and on the outer canthi of the left and right eyes. All electrode impedances were kept below 10kΩ. The EEG signal was amplified online (bandwidth: 20,000Hz) and was digitized continuously at a sampling rate of 1000Hz along with stimulus onset codes used for subsequent averaging. All channels were referenced to the right mastoid and re-referenced offline to the average of right and left mastoids. Data were collected via Pycorder (Python) and processed via Matlab (The MathWorks, Natick, MA) using ERPlab plugins (Lopez-Calderon and Luck, 2014)

*Delayed recognition test phase*. Following the EEG phase for each session, memory was tested for the words encountered during the EEG phase (64 sentence-final words, excluding those that were paired with comprehension questions during study, and 64 single words). These studied words were randomly intermixed with an equal number of novel lure words matched on length, frequency, and concreteness with studied words. Subjects attempted to discriminate studied from lure words using a four-point confidence scale (confident old, guess old, guess new, and confident new). Words were presented individually alongside the confidence scale, and subjects were given up to five seconds to respond, with the recognition response cueing delivery of the next word. The primary episodic memory outcome measure was memory performance for single words, given the uncertainty about the cognitive processes reflected by recognition of sentence-ending words.

### ERP and behavioral analysis

Analyses were performed in MATLAB (version R2017b) and R (version R-3.6.2). For ERP analyses, continuous EEG data were epoched in 1200ms intervals starting 200ms before the onset of sentence-final words and individually presented words. Independent component analysis (ICA) was performed (Li et al., 2006) to isolate and remove blink and saccade components. Components that included only frontal and ocular channels were removed (*M* = 3.8 components, *SD* = 1.5 components removed). Following ICA, single-trial waveforms were screened for artifactual contamination (i.e., muscle or non-blink eye movement) at an absolute voltage threshold of 100µV. An average of 3.1% of sentence final-word trials and 5.9% of single-word trials were excluded from further analysis (no significant differences across stimulation or day order in the number of excluded trials: *p*s > .08).

Primary analyses examined the effects of stimulation type (stimulation: cerebellar beta, cerebellar theta, location-control) on outcome measures. EEG activity during sentence-final word trials were binned by offline sentence cloze probability (cloze: high cloze, low cloze) and subsequently averaged to create ERPs. Main analyses of sentence-final words focused on the N400 (Kutas and Federmeier, 2011, Dave et al., 2018), which has typically been defined as a negative-going difference between high- and low-cloze trials maximal over central and posterior electrode sites (CP1, CP2, CZ, C3, C4, Pz, P3, P4, O1) 300-500ms after word onset. We calculated the mean ERP amplitude 300-500ms averaged for this set of electrodes for both cloze conditions and calculated the N400 by subtracting amplitudes for low cloze from those for high cloze. A repeated-measures analysis of variance (rANOVA) was performed with a within-subject factor of stimulation to assess the effect of varying stimulation parameters on N400 ERPs. All rANOVA analyses were followed by t tests of paired stimulation conditions.

We performed additional analyses on three other ERP components associated with language processing (described in Results). We measured the posterior syntactic P600 (Leckey & Federmeier, 2019) at central and posterior electrodes (CP1, CP2, CZ, C3, C4, Pz, P3, P4, O1) 600-900ms after word onset. We also calculate mean ERP amplitudes over a set of fronto-central electrodes (FP1, FP2, FC1, FC2, Fz, FC5, FC6) to assess two additional language-related effects: the N200/N250 (Holcomb and Grainger, 2006; measured for 150-300ms following word onset) and the semantic P600/PNP (Van Petten and Luka, 2012; measured for 600-900ms following word onset).

Behavioral analyses for delayed recognition of individually studied words focused on discrimination sensitivity (d′), a normalized metric of successful discrimination of studied from novel lure words (Yonelinas, 2002, Wixted, 2007). Response rates for each response type (old/new) and confidence level (confident/guess) and stimulus type (studied/foil) are listed for each condition in Table 1. As indicated in the Results, only confident responses were accurate. Therefore, we analyzed the effect of stimulation on d′ calculated only for confident responses. We performed one-way rANOVAs using a within-subjects factor of stimulation with follow-up paired t tests. We additionally examined ERP correlates of word encoding following similar procedures as for the N400 analysis. We calculated mean ERP amplitude for central and posterior electrodes where the parietal memory effect is typically maximal (CP1, CP2, CZ, C3, C4, Pz, P3, P4, O1) 650-750ms after word onset for all studied single-word trials.

## Author Contributions

S.D. and J.L.V. designed the study. S.D. collected the data. S.D. and J.L.V. performed the analyses. S.V. supervised participant eligibility and safety. All authors discussed the findings and wrote the manuscript.

## Acknowledgments

We thank M. Penckofer, M. M. Gunlogson, B. E. Durr, S. E. Lurie, and M. S. Hermiller for assistance with the experiment. Neuroimaging was performed at the Northwestern University Center for Translational Imaging, supported by the Northwestern University Department of Radiology. This research was supported, in part, through the computational resources and staff contributions provided for Quest, the high-performance computing facility at Northwestern University, which is jointly supported by the Office of the Provost, the Office for Research, and Northwestern University Information Technology.

## Funding

This research was supported by grants R01-MH106512, R01-MH111790, and F32-MH118718-01 from the National Institutes of Health. The content is solely the responsibility of the authors and does not necessarily represent the official view of the National Institutes of Health.

## Competing Interests

The authors have no competing interests to disclose.

